# Differences in *in vitro* responses of the hypothalamo-pituitary-gonadal hormonal axis between low and high egg producing turkey hens

**DOI:** 10.1101/854679

**Authors:** Kristen Brady, Julie A. Long, Hsiao-Ching Liu, Tom E. Porter

**Author notes:** Correspondence: Tom E. Porter, Ph.D., 1413 Animal Sciences Building (#142), 8127 Regents Drive, College Park, Maryland 20742. **Grant sponsor:** Agriculture and Food Research Initiative Competitive Grant #2019-67015-29472 from the USDA National Institute of Food and Agriculture.

## Abstract

Low egg producing hens (**LEPH**) ovulate less frequently than high egg producing hens (**HEPH**) and exhibit differences in mRNA levels for components of the hypothalamo-pituitary-gonadal (**HPG**) axis, suggesting differential responsiveness to trophic stimulation. Ovulation frequency is governed by the production and feedback of pituitary gonadotropins and ovarian follicle steroid hormones, which are regulated by HPG axis stimulation and inhibition at the hypothalamic level. Pituitary and follicle cells from LEPH and HEPH were subjected to *in vitro* hormonal treatments to stimulate or inhibit the HPG axis, followed by expression analysis of mRNA levels for HPG axis genes and radioimmunoassays for steroid hormone production. Statistical analysis was performed using the mixed models procedure of SAS. Pituitary cells from HEPH showed up-regulation of genes associated with ovulation stimulation, whereas LEPH cells showed up-regulation of genes associated with inhibition of ovulation. HEPH follicle cells displayed a higher sensitivity and responsiveness to gonadotropin treatment. Level of egg production impacted ovulation-related gene expression in pituitary cells as well as steroid hormone production in follicle cells, with HEPH displaying a greater positive response to stimulation. These findings indicate that differences in egg production among turkey hens likely involve differential responsiveness of the cells within the HPG axis.

## INTRODUCTION

Differences in egg production rates among turkey hens in a flock result in low egg producing hens (**LEPH**) and high egg producing hens (**HEPH**). Low egg production in breeding hens costs the industry in lost poult production and is correlated with decreased ovulation frequency (Liu et al., 2005). Follicle ovulation in avian species is controlled by the hypothalamo-pituitary-gonadal (**HPG**) axis, which is composed of the hypothalamus, pituitary, and a single ovary. A preovulatory surge (**PS**) precedes each ovulation and consists of increased progesterone and luteinizing hormone (**LH**), produced by the ovary and pituitary, respectively (Paster, 1991). Steroid hormones, estradiol and progesterone, feedback on the HPG axis to regulate ovulation timing (Ottinger and Bakst, 1995).

The HPG axis can be stimulated by gonadotropin releasing hormone (**GnRH**) or inhibited by gonadotropin inhibitory hormone (**GnIH**), both produced in the hypothalamus with the anterior pituitary as their target tissue. Neuron terminals containing GnRH extend into the external layer of the median eminence for neuropeptide release into the hypophysial portal vascular system (Bédécarrats, 2014). Neuron terminals containing GnIH also extend into the median eminence but also have direct contact with GnRH neurons, suggesting the capability of GnIH regulation of GnRH synthesis and release (Bédécarrats et al., 2016). GnRH or GnIH regulate pituitary gonadotropin production by binding to either gonadotropin releasing hormone receptor (**GnRHR**) or gonadotropin inhibitory hormone receptor (**GnIHR**), both located on pituitary gonadotrophs. GnRHR and GnIHR are G protein-coupled receptors (**GPCRs**) present on pituitary gonadotroph cells, with GnRHR coupling to G_αs_ and G_αq_ and GnIHR coupling to G_αi_ (Tsutsui et al., 2006).

The ovary is composed of follicles in varying states of maturation, developing from quiescent primordial follicles to preovulatory follicles awaiting ovulation. Steroidogenesis occurs in ovarian follicles, with primary steroid production varying with follicle development and follicle cell type (Porter et al., 1989). The majority of ovarian estradiol production occurs in the small white follicles (**SWF**), which are slow growing follicles that have yet to enter the preovulatory hierarchy (Johnson, 1992). Ovarian progesterone production primarily occurs in the granulosa cells of the largest preovulatory follicle (**F1G**), which is the next follicle in line to ovulate (Bahr et al., 2005). SWF are mainly responsive to follicle stimulating hormone (**FSH**), while F1G are responsive to LH. FSH receptor (**FSHR**) and LH receptor (**LHCGR**) are also GPCRs that couple to G_αs_ to increase the transcription of genes involved in steroidogenesis through the cAMP-signaling pathway, such as steroidogenic acute regulatory protein (***STAR***) and aromatase (***CYP19A1***) (Li et al., 2014).

Previous studies comparing HPG axis gene expression of LEPH and HEPH found that HEPH displayed gene expression levels consistent with increased ovulation stimulation and decreased ovulation inhibition in the hypothalamus and pituitary. Additionally, HEPH showed upregulation of genes related to progesterone production in the F1G and related to estradiol production in SWF (Brady et al., 2019a). HEPH showed decreased gene expression of *GNIH* in the hypothalamus, increased expression of both gonadotropin beta-subunits in the pituitary, increased gene expression of *STAR* and cholesterol side chain cleavage enzyme (***CYP11A1***) in the F1G, and increased gene expression of 17β-hydroxysteroid dehydrogenase (***HSD17B1***) and *CYP19A1* in the SWF.

Based on previous gene expression differences between LEPH and HEPH, it was hypothesized that HEPH would show an increased sensitivity and responsiveness to GnRH treatment in the pituitary, to LH treatment in the F1G, and to FSH treatment in the SWF, whereas LEPH would show an increased sensitivity and responsiveness to GnIH treatment in the pituitary. This study sought to compare LEPH and HEPH in the responsiveness of isolated pituitary cells to GnRH and GnIH stimulation as well as in the responsiveness of the two follicle cell types responsible for estradiol and progesterone production, SWF and F1G, to FSH and LH stimulation, respectively.

## MATERIALS AND METHODS

### Hen Selection

Females (200 hens) from a commercial line (Hybrid Turkey, Kitchener, Ontario) were housed at the Beltsville Agricultural Research Center (**BARC**) in individual wire cages. Turkey hens were maintained under standard poultry management practices with artificial lighting (14L:10D) and were provided feed *ad libitum* to NRC standards. Hens were sampled at 35 weeks of age. Daily egg records were used to calculate each hen’s number of eggs per day (**EPD**) by dividing the total number of eggs produced by the number of days in production. Hens were classified as LEPH when EPD<0.6 and as HEPH when EPD>0.8. Egg production cutoffs were based on the distribution of flock egg production as previously described (Brady et al., 2019a). Blood samples were taken from the wing vein immediately before sampling, collected in heparinized tubes, and fractionated by centrifugation (2000xg for 10 minutes at room temperature). Plasma samples were stored at −20°C prior to assessment through radioimmunoassays as described below. The pituitary, F1 follicle, and SWF were isolated from four LEPH and four HEPH. All hens were sampled at the same time during the daily lighting schedule, sampled outside of the PS, and sampled on the second day of the hen’s sequence, as previously described (Brady et al., 2019b). The timing of the PS was estimated for each hen based on the oviposition-ovulation cycle as previously described (Brady et al., 2019b). Plasma progesterone levels were examined to confirm correct sampling time during the ovulatory cycle. All animal procedures were approved by the Institutional Animal Care and Use Committee at BARC and at the University of Maryland.

### Cell Isolation and Culture

All cell isolation procedures were performed using Minimum Essential Medium, Spinner modification (SMEM) or Dulbecco’s Modified Eagle Medium (DMEM) as noted below. Media was supplemented with 0.1% bovine serum albumen, 100-U/mL penicillin G, and 100-μg/mL streptomycin sulfate (0.1% BSA and P/S).

Isolated pituitaries were minced and dispersed in SMEM (0.1% BSA and P/S) using trypsin and collagenase (1 mg/mL of each) for 90 minutes at 37°C in a shaking water bath. After dispersion, cells were filtered through 70 μm nylon mesh and washed twice with DMEM (0.1% BSA and P/S). Cells were diluted to a concentration of 200,000 cells/mL and plated in serum free medium (DMEM/F12) supplemented with 0.1% bovine serum albumen, 5-μg/mL human insulin, 100-U/mL penicillin G, and 100-μg/mL streptomycin sulfate. Cells were plated in 24-well poly-L lysine coated plates (Corning Life Sciences, Lowell, MA) at 100,000 cells/well and were allowed to attach for 2 hours before treatment. Pituitary cells were treated with chicken GnRH or GnIH (Phoenix Pharmaceuticals, Burlingame, CA) at 0, 10^-9^, 10^-8^, or 10^-7^ M for 6 or 24 hours. Timepoints were selected to examine short- and long-term gene expression changes due to treatment.

The F1 follicle was removed from the ovary and placed in ice cold SMEM (0.1% BSA and P/S) until isolation of the granulosa cell layer. The follicle was drained of yolk, everted, and the granulosa cell layer was peeled off of the follicle wall. The granulosa cell layer was dispersed in SMEM (0.1% BSA and P/S) using trypsin (1 mg/mL) as previously described (Brady et al., 2019b). After dispersion, cells were filtered through 70 μm nylon mesh and layered on a 50% Percoll solution to remove remaining yolk particles. Cells were washed twice with SMEM (0.1% BSA and P/S) and diluted to a density of 10,000 cells/mL for culture. F1G cells were cultured in SMEM (0.1% BSA and P/S) in 12x75 mm polypropylene tubes (1x10^5^ cells per tube). Cells were treated with ovine LH (National Hormone & Peptide Program, Torrance, CA) at 0, 1, 10, 100, or 1000 ng/mL for 5 hours as previously described (Porter et al., 1989).

SWF were minced and dispersed in SMEM (0.1% BSA and P/S) using trypsin (1 mg/mL) for 60 minutes at 37°C in a shaking water bath. After dispersion, cells were filtered through 70 μm nylon mesh and layered on a 50% Percoll solution to remove remaining red blood cells. SWF cells were washed twice with SMEM (0.1% BSA and P/S) and diluted to a density of 10,000 cells/mL for culture. SWF cells were cultured in SMEM (0.1% BSA and P/S) in 12x75 mm polypropylene tubes (1x10^5^ cells per tube). Cells were treated with porcine FSH (National Hormone & Peptide Program, Torrance, CA) at 0, 1, 10, 100, or 1000 ng/mL for 5 hours as previously described (Porter et al., 1989).

Cells were maintained in a 37.5°C, 5% CO_2_ atmosphere during incubation. Pituitary cells were harvested at the completion of each incubation by retrypsinization, immediately frozen in liquid nitrogen, and stored at −80 °C until RNA extraction. The media from the F1G and SWF cell cultures was recovered and stored at −20 °C for progesterone and estradiol radioimmunoassays, respectively.

### RT-qPCR

Total RNA was isolated from pituitary cell cultures with RNeasy Mini kits (Qiagen, Valencia, CA), including on-column deoxyribonuclease digestion. Quantification of RNA, RT, and RT-qPCR were performed as previously described (Brady et al., 2019b) with the following exception. Reverse transcription reactions were performed on 50 ng total RNA with SuperScript III (Thermo Fisher Scientific, Waltham, MA) and an anchored oligo-dT primer (5′-CGGAATTCTTTTTTTTTTTTTTTTTTTTV-3′) (Integrated DNA Technologies, Skokie, IL). A pool of total RNA was made and the reaction conducted without reverse transcriptase as a control for genomic DNA contamination. Reactions were diluted to 40 μl prior to PCR analysis. PCR reactions (15 μL) were carried out as previously described using a CFX Connect Real-Time PCR System (Bio-Rad, Hercules, CA) (Brady et al., 2019b). Data were normalized to phosphoglycerate kinase 1 (*PGK1*) and analyzed by the 2^-ΔΔ Ct^ method. All PCR reactions for each gene in a given tissue were analyzed in a single run within a 96-well plate, allowing accurate performance of relative quantification without the need to include a reference control sample in each plate. Primers (Integrated DNA Technologies, Skokie, IL) for turkey *PGK1*, gonadotropin releasing hormone receptor (*GNRHR*), gonadotropin inhibitory hormone receptor (*GNIHR*), luteinizing hormone beta-subunit (*LHB*), follicle stimulating hormone beta-subunit (*FSHB*), and glycoprotein hormone alpha-subunit (*CGA*) mRNA were designed and used with cycling parameters described previously (Brady et al., 2019b). Data are presented as fold increase over levels in basal cells for each hormone treatment and time point.

### Radioimmunoassay (RIA)

The RIAs used for progesterone and estradiol were coated tube kits (MP Biomedicals, Solon, OH). All protocols were performed as directed by the supplier. All samples were assayed in duplicate. All samples were measured in a single RIA for each hormone. Plasma samples were ether extracted and analyzed for progesterone to determine that hens were sampled outside of the preovulatory surge. Culture media from the F1G and SWF cell cultures were assayed for progesterone and estradiol content, respectively. The standard curve was assessed for linearity as well as dilutional parallelism using serial plasma or culture media dilutions. The intraassay coefficients of variation determined by pools run every 30 samples were 5.61% for progesterone and 6.63% for estradiol.

### Statistics

All data were analyzed using SAS software (SAS Institute, Cary, NC). Normalized RT-qPCR data were log_2_ transformed before statistical analysis. A three-way ANOVA using the mixed models procedure (PROC MIXED) was conducted to compare the log_2_ transformed pituitary gene expression between LEPH and HEPH. A two-way ANOVA using the mixed models procedure (PROC MIXED) was used to compare plasma hormone concentrations and culture media hormone concentrations between LEPH and HEPH. The least squares means for each group were compared using the test of least significant difference (PDIFF statement) when this indicated an overall significance level of *P* < 0.05.

## RESULTS

*GNRHR* expression in response to GnRH treatment is presented in **Figure 1a**. After GnRH treatment for 6 hours, pituitary cells from HEPH showed higher *GNRHR* expression relative to cells from LEPH at 10^-9^ and 10^-8^ M GnRH, while cells from LEPH showed higher *GNRHR* expression relative to cells from HEPH at 10^-7^ M GnRH. Cells from LEPH showed increased *GNRHR* expression relative to basal expression in response to GnRH treatment only at 10^-7^ M. Cells from HEPH showed increased *GNRHR* expression relative to basal expression in response to GnRH treatment only at 10^-8^ M. *GNRHR* expression was not affected by GnRH treatment for 24 hours.

**Figure 1.**
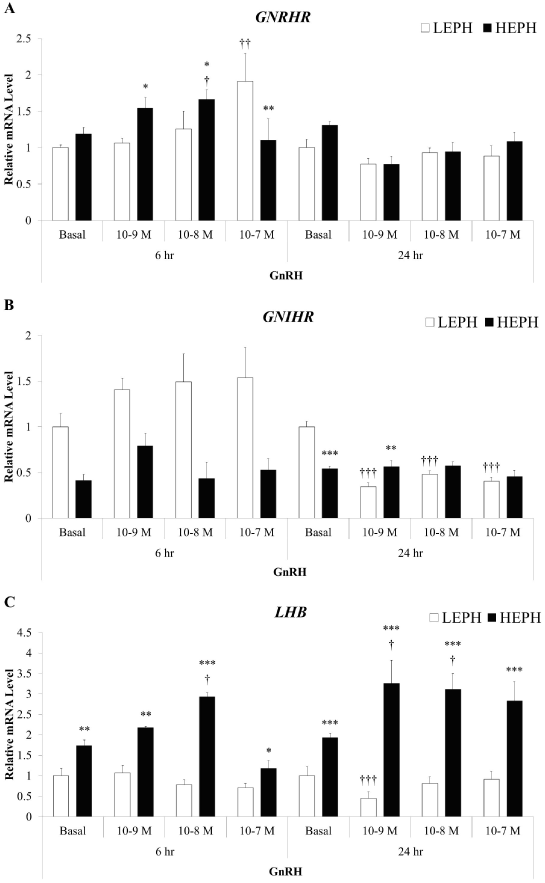
Relative pituitary expression of gonadotropin-releasing hormone receptor (*GNRHR*), gonadotropin-inhibitory hormone receptor (*GNIHR*) and the beta subunit of luteinizing hormone (*LHB*) after gonadotropin-releasing hormone (GNRH) treatment in low egg producing hens (LEPH) and high egg producing hens (HEPH). Normalized data are presented relative to LEPH basal expression for each gene. Significant expression differences between LEPH and HEPH for a given condition are denoted with an asterisk (*P≤0.05, **P≤0.01, and ***P≤0.001), whereas, significant differences between basal and a specific GnRH treatment for a given egg production group are denoted with a dagger (†P≤0.05, ††P≤0.01, and †††P≤0.001).

*GNIHR* expression in response to GnRH treatment is presented in **Figure 1b**. After GnRH treatment for 6 hours, *GNIHR* expression was significantly affected by egg production level but a response to GnRH treatment was not seen. After GnRH treatment for 24 hours, pituitary cells from LEPH showed higher *GNIHR* expression relative to cells from HEPH under basal conditions, but HEPH demonstrated higher *GNIHR* expression at 10^-9^ M GnRH. Cells from LEPH showed decreased *GNIHR* expression relative to basal expression in response to all GnRH treatments. Cells from HEPH did not show a response in *GNIHR* expression to GnRH treatment.

*LHB* expression in response to GnRH treatment is presented in **Figure 1c**. After GnRH treatment for 6 hours, pituitary cells from HEPH showed higher *LHB* expression relative to cells from LEPH at all GnRH treatments. Cells from LEPH did not show a response in *LHB* expression after GnRH treatment. Cells from HEPH showed increased *LHB* expression relative to basal expression in response to GnRH treatment at 10^-8^ M and decreased expression relative to basal expression at 10^-7^ M GnRH. After GnRH treatment for 24 hours, pituitary cells from HEPH showed higher *LHB* expression relative to cells from LEPH at all GnRH treatments. Cells from LEPH showed decreased *LHB* expression after GnRH treatment at 10^-9^ M. Cells from HEPH showed increased *LHB* expression relative to basal expression in response to GnRH treatment at 10^-9^ and 10^-8^ M. *GNRHR* expression in response to GnIH treatment is presented in **Figure 2a**. After GnIH treatment for 6 hours, pituitary cells from HEPH showed higher *GNRHR* expression relative to cells from LEPH at 10^-9^ M GnIH, while cells from LEPH showed higher *GNRHR* expression relative to cells from HEPH at 10^-7^ M GnIH. Cells from LEPH showed increased *GNRHR* expression relative to basal expression in response to GnIH treatment at 10^-8^ and 10^-7^ M. Cells from HEPH showed increased *GNRHR* expression relative to basal expression in response to GnIH treatment at 10^-9^ and 10^-8^ M before returning to basal expression. After GnIH treatment for 24 hours, pituitary cells from HEPH showed higher *GNRHR* expression relative to cells from LEPH under basal conditions while cells from LEPH showed higher *GNRHR* expression relative to cells from HEPH at 10^-7^ M GnIH. Cells from LEPH showed increased *GNRHR* expression relative to basal expression in response to all the GnIH concentrations, while cells from HEPH did not show changes in *GNRHR* expression across GnIH treatments.

**Figure 2.**
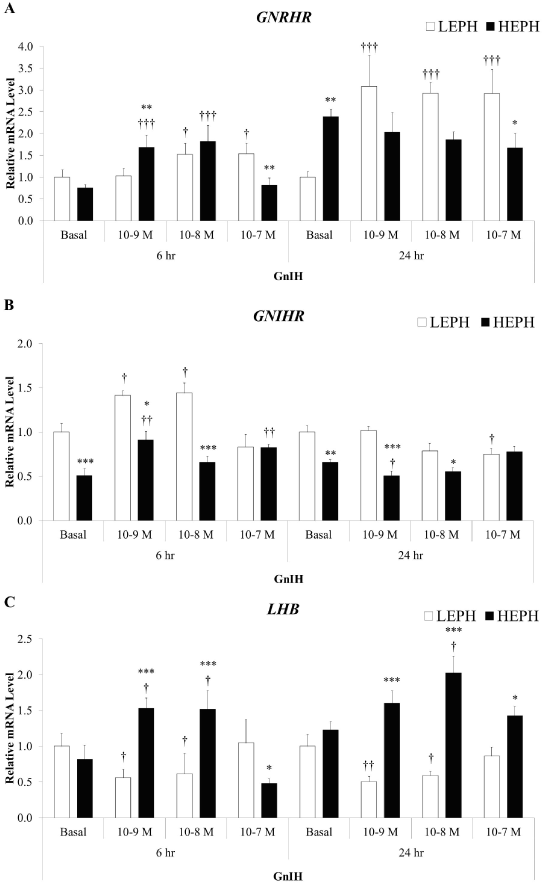
Relative pituitary expression of gonadotropin-releasing hormone receptor (*GNRHR*), gonadotropin-inhibitory hormone receptor (*GNIHR*) and the beta subunit of luteinizing hormone (*LHB*) after gonadotropin-inhibitory hormone (GNIH) treatment in low egg producing hens (LEPH) and high egg producing hens (HEPH). Normalized data are presented relative to LEPH basal expression for each gene. Significant expression differences between LEPH and HEPH for a given condition are denoted with an asterisk (*P≤0.05, **P≤0.01, and ***P≤0.001), whereas, significant differences between basal and a specific GnRH treatment for a given egg production group are denoted with a dagger (†P≤0.05, ††P≤0.01, and †††P≤0.001).

*GNIHR* expression in response to GnIH treatment is presented in **Figure 2b**. After GnIH treatment for 6 hours, pituitary cells from LEPH showed higher *GNIHR* expression relative to cells from HEPH at 0, 10^-9^, and 10^-8^ M GnIH. Cells from LEPH showed increased *GNIHR* expression relative to basal expression in response to GnIH treatment at 10^-9^ and 10^-8^ M before returning to basal expression. Cells from HEPH showed increased *GNIHR* expression relative to basal expression in response to GnIH treatment at 10^-9^ and 10^-7^ M. After GnIH treatment for 24 hours, pituitary cells from LEPH showed higher *GNIHR* expression relative to cells from HEPH at 0, 10^-9^, and 10^-8^ M GnIH. Cells from LEPH showed decreased *GNIHR* expression relative to basal expression in response to GnIH treatment at 10^-7^ M. Cells from HEPH showed decreased *GNIHR* expression relative to basal expression before returning to basal expression in response to GnIH treatment at 10^-9^ M.

*LHB* expression in response to GnIH treatment is presented in **Figure 2c**. After GnIH treatment for 6 hours, pituitary cells from HEPH showed higher *LHB* expression relative to cells from LEPH at 10^-9^ and 10^-8^ M GnIH, while LEPH showed higher *LHB* expression at 10^-7^ M GnIH. Cells from LEPH showed decreased *LHB* expression relative to basal expression in response to GnIH treatment at 10^-9^ and 10^-8^ M before returning to basal expression. Cells from HEPH showed increased *LHB* expression relative to basal expression in response to GnIH treatment at 10^-9^ and 10^-8^ M before reducing expression at 10^-7^ M GnIH. After GnIH treatment for 24 hours, pituitary cells from HEPH showed increased *LHB* expression relative to cells from LEPH at 10^-9^, 10^-8^, and 10^-7^ M GnIH. Cells from LEPH showed decreased *LHB* expression relative to basal expression in response to GnIH treatment at 10^-9^ and 10^-8^ M before returning to basal expression. Cells from HEPH showed increased *LHB* expression relative to basal expression in response to all GnIH treatments.

*FSHB* and *CGA* expression was not affected by GnRH or GnIH treatment for 6 or 24 hours, and results are not presented.

F1G cell progesterone production in response to LH treatment is presented in **Figure 3**. Basal progesterone production from F1G cells from LEPH and HEPH did not differ significantly. F1G cells from HEPH responded to LH treatment with increased progesterone production compared to basal levels at 10, 100, and 1000 ng/mL, whereas F1G cells from LEPH did not respond to LH treatment treatment with increased progesterone production compared to basal levels at the four experimental concentrations. Progesterone production differed significantly between F1G cells from LEPH and HEPH after treatment with 10, 100, and 1000 ng/mL of LH.

**Figure 3.**
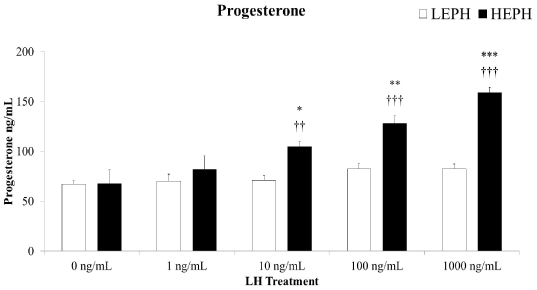
Progesterone production in F1 follicle granulosa cells (F1G) from low egg producing hens (LEPH) and high egg producing hens (HEPH) after luteinizing hormone (LH) treatment. Significant expression differences between LEPH and HEPH for a given condition are denoted with an asterisk (*P≤0.05, **P≤0.01, and ***P≤0.001), whereas, significant differences between basal and a specific LH treatment for a given egg production group are denoted with a dagger (†P≤0.05, ††P≤0.01, and †††P≤0.001).

SWF cell estradiol production in response to FSH treatment is presented in **Figure 4**. Basal estradiol production from SWF cells from LEPH and HEPH did not differ significantly. SWF cells from HEPH responded to FSH treatment with increased estradiol production compared to basal levels at 10, 100, and 1000 ng/mL, whereas SWF cells from LEPH only responded to FSH treatment with increased estradiol production compared to basal levels at 100 and 1000 ng/mL. Estradiol production differed significantly between SWF cells from LEPH and HEPH after treatment with 10, 100, and 1000 ng/mL of FSH.

**Figure 4.**
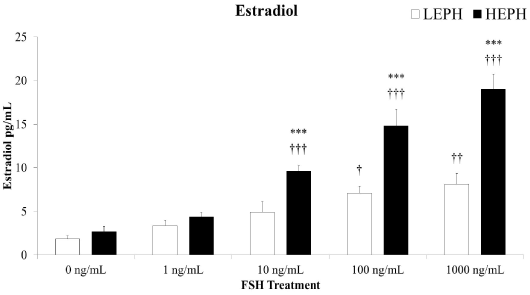
Estradiol production in small white follicle cells (SWF) from low egg producing hens (LEPH) and high egg producing hens (HEPH) after follicle stimulating hormone (FSH) treatment Significant expression differences between LEPH and HEPH for a given condition are denoted with an asterisk (*P≤0.05, **P≤0.01, and ***P≤0.001), whereas, significant differences between basal and a specific FSH treatment for a given egg production group are denoted with a dagger (†P≤0.05, ††P≤0.01, and †††P≤0.001).

## DISCUSSION

The current study showed that pituitary, F1G, and SWF cells from LEPH and HEPH respond differently to HPG axis hormone stimulation and inhibition. Previous studies have focused on HPG axis hormone responses in pituitary and ovarian cells during the initiation of egg production or during gonadal regression in both chicken and turkey hens (Guémené and Williams, 1999; Porter et al., 1991). Additionally, previous studies have compared ovarian response to gonadotropin stimulation in broiler and layer line chicken hens (Hocking and McCormack, 2004). This is the first study to examine HPG axis hormone response differences in hens with differential egg production from the same flock. Differences in *in vitro* responses to stimulation and inhibition coupled with previously identified HPG axis gene expression differences suggest core differences in the regulation of HPG axis function between LEPH and HEPH.

GnRH treatment positively impacted the expression of *GNRHR*, *GNIHR*, and *LHB* in the pituitary cells from HEPH. Receptor gene expression changes occurred during short-term GnRH treatment, while *LHB* gene expression changes were seen during short-term and long-term GnRH treatment. Cells from LEPH displayed minimal increased expression of *GNRHR*, decreased expression of *GNIHR*, and no changes in expression of *LHB* in response to GnRH treatment. Expression of *GNRHR* and *GNIHR* after GnRH treatment has not been previously examined in avian species; however, studies in a mammalian gonadotroph cell line showed up-regulation of *GNRHR* and *GNIHR* following GnRH treatment (Turgeon et al., 2014; Sukhbaatar et al., 2014). Injection of GnRH in chickens resulted in increased plasma LH in previous studies (Wilson et al., 1989). While the results from the current study examine the transcriptional changes due to GnRH treatment, increased *LHB* expression is consistent with studies at the protein level. *LHB* expression decreased to basal levels at the highest short-term GnRH treatment. Similar desensitization has been shown in prior studies (King et al., 1986). *FSHB* expression was not affected by GnRH treatment in LEPH or HEPH, which is consistent with prior studies (Proudman et al., 2006). Increased responsiveness to GnRH in HEPH, in terms of *LHB* and *GNRHR* expression, may result in increased ovulation rates in these hens, through increased gonadotropin production.

While GnIH treatment up-regulated *GNRHR* expression in both groups of hens, *GNIHR* expression was only up-regulated in cells from LEPH, and *LHB* expression was only up-regulated in cells from HEPH following GnIH treatment. *In vitro*, activation of *GNIHR* through GnIH binding has been shown to reduce *GNRHR* gene expression in chicken pituitary cells (Bédécarrats et al., 2009). Cells from LEPH and HEPH initially displayed up-regulation of *GNRHR* after GnIH treatment, though long-term treatment resulted in a further increase in *GNRHR* expression in cells from LEPH and in mRNA levels similar to basal expression in cells from HEPH. GnIH treatment has been shown to increase *GNIHR* expression in the chicken (Maddineni et al., 2008). Cells from LEPH and HEPH both showed short-term up-regulation of *GNIHR* after GnIH treatment. However, cells from LEPH displayed higher overall mRNA levels of *GNIHR* for each GnIH treatment. GnIH has also been shown to decrease LH synthesis and release in the chicken, both *in vivo* and *in vitro* (Bédécarrats et al., 2009). This phenomenon was seen in cells from LEPH after GnIH treatment but up-regulation in response to GnIH treatment was seen in cells from HEPH. This may indicate that GnIH signaling in LEPH and HEPH operates differently to control gonadotropin production. Studies in mammalian gonadotroph cell lines indicated that GnIH treatment decreased the expression of *FSHB* and *CGA*, both of which were not seen in the current study (Son et al., 2012). Differences in GnIH regulation may be attributed to the reproductive physiology differences between mammalian and avian species. Increased responsiveness of LEPH to GnIH through increased *GNIHR* expression may allow for the up-regulation of HPG axis inhibitory pathways, decreasing gonadotropin production.

A dose-dependent progesterone production response to LH treatment was seen in F1G cells from HEPH, whereas cells from LEPH did not respond to LH treatment in terms of progesterone production. Results seen in HEPH cells are consistent with previously published studies in chicken and turkey hens (Bakst et al., 1983; Porter et al., 1991). Progesterone production from the F1G cells is imperative for the PS to occur and induced ovulation. Lack of response to LH stimulation in F1G cells from LEPH may contribute to decreased ovulation rates seen in this group of hens. An estradiol production response to FSH treatment was seen in SWF cells from LEPH and HEPH; however, SWF cells from HEPH responded at a lower dose of FSH treatment and responded with significantly higher estradiol production when compared to cells from LEPH. The SWF are the follicle cell type with the greatest FSH binding. Results seen in HEPH were also consistent with previous studies in chicken and turkey SWF (Etches and Cheng, 1981; Porter et al., 1989). Furthermore, F1G and SWF results from the current study are consistent with previous results showing up-regulation of genes in cells from HEPH when compared to cells from LEPH that are involved in progesterone and estradiol production (Brady et al., 2019^a^). Follicles from LEPH and HEPH appear to respond differently to gonadotropin stimulation, impacting the steroid hormone production capabilities in these groups of hens. Increased progesterone and estradiol production in HEPH may lead to increased egg production levels through local ovarian effects decreasing ovulation intervals or through HPG axis feedback at the hypothalamic and pituitary level regulating releasing factor and gonadotropin production.

In summary, HEPH displayed increased responsiveness to GnRH in pituitary cells, to LH in F1G cells, and to FSH in SWF cells. On the other hand, pituitary cells from LEPH displayed increased responsiveness to the inhibitory properties of GnIH, whereas pituitary cells from HEPH responded positively to GnIH treatment. These findings demonstrate that HPG axis responsiveness is different in LEPH and HEPH, with LEPH favoring the inhibitory pathways of the axis and HEPH favoring the stimulatory pathways of the axis. Understanding how HPG axis hormone responsiveness ultimately impacts egg production on a molecular level would be imperative to improve the reproductive efficiency of LEPH in the turkey industry.

## ACKNOWLEDGEMENTS

This project was supported by Agriculture and Food Research Initiative Competitive Grant no. 2019-67015-29472 from the USDA National Institute of Food and Agriculture.

## REFERENCES

1. Bahr, J. M., Wang, S.-C., Huang, M. Y., & Calvo, F. O. (1983). Steroid Concentrations in Isolated Theca and Granulosa Layers of Preovulatory Follicles During the Ovulatory Cycle of the Domestic Hen. Biology of Reproduction, 29, 326–334. *doi*:10.1095/biolreprod29.2.326

2. Bakst, M., Scanes, C., & Phillips, A. (1983). Steroidogenic and morphological responses of hen granulosa cells to luteinizing hormone *in vitro*. *Scanning Electron Microscopy*, (Pt 4), 1931–1938.

3. Brady, K., Porter, T. E., Liu, H. C., & Long, J. A. (2019). Characterization of the hypothalamo-pituitary-gonadal axis in low and high egg producing turkey hens. *Poultry Science*, pez579. *doi*:10.3382/ps/pez579

4. Brady, K., Porter, T. E., Liu, H. C., & Long, J. A. (2019). Characterization of gene expression in the hypothalamo-pituitary-gonadal axis during the preovulatory surge in the turkey hen. *Poultry Science*, pez437. *doi*:10.3382/ps/pez437

5. Bédécarrats, G. Y. (2014). Control of the reproductive axis: Balancing act between stimulatory and inhibitory inputs. Poultry Science, 94, 810–815. *doi*:10.3382/ps/peu042

6. Bédécarrats, G. Y., Baxter, M., & Sparling, B. (2016). An updated model to describe the neuroendocrine control of reproduction in chickens. General and Comparative Endocrinology, 227, 58–63. *doi*:10.1016/j.ygcen.2015.09.023

7. Bédécarrats, G. Y., McFarlane, H., Maddineni, S. R., & Ramachandran, R. (2009). Gonadotropin-inhibitory hormone receptor signaling and its impact on reproduction in chickens. General and Comparative Endocrinology, 163, 7–11. *doi*:10.1016/j.ygcen.2009.03.010

8. Etches, R. J., & Cheng, K. W. (1981). Changes in the plasma concentrations of luteinizing hormone, progesterone, oestradiol and testosterone and in the binding of follicle-stimulating hormone to the theca of follicles during the ovulation cycle of the hen (Gallus domesticus). Journal of Endocrinology, 91, 11–22. *doi*:10.1677/joe.0.0910011

9. Guémené, D., & Williams, J. B. (1999). LH responses to chicken luteinizing hormone-releasing hormone I and II in laying, incubating, and out of lay turkey hens. Domestic Animal Endocrinology, 17, 1–15. *doi*:10.1016/S0739-7240(99)00020-X

10. Hocking, P. M., & McCormack, H. A. (1995). Differential sensitivity of ovarian follicles to gonadotrophin stimulation in broiler and layer lines of domestic fowl. Journal of Reproduction and Fertility, 105, 49–55. *doi*:10.1530/jrf.0.1050049

11. Johnson, A. L. (1992). Some Characteristics of Small White Ovarian Follicles: Implications for Recruitment and Atresia in the Domestic Hen. Scandinavian Journal of Ornithology, 23, 233–237. *doi*:10.2307/3676643

12. King, J. A., Davidson, J. S., & Millar, R. P. (1986). Desensitization to gonadotropin-releasing hormone in perifused chicken anterior pituitary cells. Endocrinology, 119, 1510–1518. *doi*:10.1210/endo-119-4-1510

13. Li, L., Zhang, Z., Peng, J., Wang, Y., & Zhu, Q. (2014). Cooperation of luteinizing hormone signaling pathways in preovulatory avian follicles regulates circadian clock expression in granulosa cell. Molecular and Cellular Biochemistry, 394, 31–41. *doi*:10.1007/s11010-014-2078-3

14. Liu, H.-K., Nestor, K. E., Long, D. W., & Bacon, W. L. (2001). Frequency of Luteinizing Hormone Surges and Egg Production Rate in Turkey Hens. Biology of Reproduction, 64, 1769–1775. *doi*:10.1095/biolreprod64.6.1769

15. Maddineni, S., Ocón-grove, O. M., Krzysik-Walker, S. M., Hendricks, G. L., Proudman, J. A., & Ramachandran, R. (2008). Gonadotrophin-Inhibitory hormone receptor expression in the chicken pituitary gland: Potential influence of sexual maturation and ovarian steroids. Journal of Neuroendocrinology, 20, 1078–1088. *doi*:10.1111/j.1365-2826.2008.01765.x

16. Ottinger, Mary, Bakst, M. (1995). Endocrinology of the Avian Reproductive System. Journal of Avian Medicine and Surgery, 9, 242–250.

17. Paster, M. B. (1991). Avian reproductive endocrinology. The Veterinary Clinics of North America Small Animal Practice, 21, 1343–1359. *doi*:10.1016/S0195-5616(91)50143-1

18. Porter, T. E., Hargis, B. M., Silsby, J. L., & El Halawani, M. E. (1989). Enhanced progesterone and testosterone secretion and depressed estradiol secretion in vitro from small white follicle cells of incubating turkey hens. General and Comparative Endocrinology, 74, 400–405. *doi*:10.1016/S0016-6480(89)80037-1

19. Porter, T. E., Hargis, B. M., Silsby, J. L., & Halawani, M. E. E. (1989). Differential steroid production between theca interna and theca externa cells: A three-cell model for follicular steroidogenesis in avian species. Endocrinology, 125, 109– 116. *doi*:10.1210/endo-125-1-109

20. Porter, T. E., Hargis, B. M., Silsby, J. L., & El Halawani, M. E. (1991). Characterization of dissimilar steroid productions by granulosa, theca interna and theca externa cells during follicular maturation in the turkey (Meleagris gallopavo). General and Comparative Endocrinology, 84, 1–8. *doi*:10.1016/0016-6480(91)90058-E

21. Porter, T. E., Silsby, J. L., Behnke, E. J., Knapp, T. R., & El Halawani, M. E. (1991). Ovarian steroid production in vitro during gonadal regression in the turkey. I. Changes associated with incubation behavior. Biology of Reproduction, 45, 581–586. *doi*:10.1095/biolreprod45.4.581

22. Proudman, J. A., Scanes, C. G., Johannsen, S. A., Berghman, L. R., & Camp, M. J. (2006). Comparison of the ability of the three endogenous GnRHs to stimulate release of follicle-stimulating hormone and luteinizing hormone in chickens. Domestic Animal Endocrinology, 31, 141–153. *doi*:10.1016/j.domaniend.2005.10.002

23. Son, Y. L., Ubuka, T., Millar, R. P., Kanasaki, H., & Tsutsui, K. (2012). Gonadotropin-inhibitory hormone inhibits gnrh-induced gonadotropin subunit gene transcriptions by inhibiting AC/cAMP/PKA-dependent ERK pathway in LβT2 Cells. Endocrinology, 153, 2332–2343. *doi*:10.1210/en.2011-1904

24. Sukhbaatar, U., Kanasaki, H., Mijiddorj, T., Oride, A., & Miyazaki, K. (2014). Expression of gonadotropin-inhibitory hormone receptors in mouse pituitary gonadotroph LβT2 cells and hypothalamic. Endocrine Journal, 61, 25–34. *doi*:10.1507/endocrj.ej13-0238

25. Tsutsui, K., Ubuka, T., Yin, H., Osugi, T., Ukena, K., Bentley, G. E., … Wingfield, J. C. (2006). Mode of action and functional significance of avian gonadotropin-inhibitory hormone (GnIH): A review. Journal of Experimental Zoology Part A: Comparative Experimental Biology, 305, 801–806. *doi*:10.1002/jez.a.305

26. Turgeon, J. L., Kimura, Y., Waring, D. W., & Mellon, P. L. (1996). Steroid and pulsatile gonadotropin-releasing hormone (GnRH) regulation of luteinizing hormone and GnRH receptor in a novel gonadotrope cell line. Molecular Endocrinology, 10, 439–450. *doi*:10.1210/mend.10.4.8721988

27. Wilson, S. C., Cunningham, F. J., Chairil, R. A., & Gladwell, R. T. (1989). Maturational changes in the LH response of domestic fowl to synthetic chicken LHRH-I and -II. Journal of Endocrinology, 123, 311–318. *doi*:10.1677/joe.0.1230311

